# A missense mutation in *Muc2* promotes gut microbiome- and metabolome-dependent colitis-associated tumorigenesis

**DOI:** 10.1101/2025.05.31.657160

**Authors:** Giulio Verna, Stefania De Santis, Bianca Islam, Eduardo M. Sommella, Danilo Licastro, Liangliang Zhang, Fabiano De Almeida Celio, Fabrizio Merciai, Vicky Caponigro, Pietro Campiglia, Theresa T. Pizarro, Marcello Chieppa, Fabio Cominelli

## Abstract

Colitis-associated cancer (CAC) arises from a complex interplay between host and environmental factors, including the gut microbiome. Since ulcerative colitis (UC), a significant risk factor for CAC, is rising in prevalence worldwide, an integrative approach is essential to identify potential triggers linking inflammation to cancer. In the present study, we investigated the role of the gut microbiome using Winnie mice, a UC-like model with a relevant missense mutation in the *Muc2 gene.* Upon transfer from a conventional (CONV) to a specific-pathogen-free (SPF) facility, Winnie mice exhibited a more severe colitis phenotype, and notably, spontaneous CAC as early as four weeks of age, which progressively worsened over time. In contrast, CONV Winnie developed only mild colitis but with no overt signs of tumorigenesis. Notably, when rederived into germ-free (GF) conditions, SPF Winnie mice were protected from colitis or colon tumor development, indicating an essential role for the gut microbiome in the initiation and progression of CAC in these mice. Using shotgun metagenomics, metabolomics, and lipidomics, we identified a distinct pro-inflammatory microbial and metabolic signature that potentially drives the transition from colitis to CAC. Fecal microbiota transplantation (FMT), using either SPF Winnie or WT (Bl/6) donors into GF Winnie recipients, demonstrated that while colitis developed regardless of donor, only FMT from SPF Winnie donors resulted in CAC, revealing a microbiota-driven, host-specific susceptibility to tumorigenesis in Winnie mice. Our studies present a novel and relevant model of CAC, providing further evidence that the microbiome plays a key role in the pathogenesis of CAC, thereby challenging the concept of colon cancer as a strictly non-transmissible disease.

**Lay summary:** This study reveals a distinct metagenomic, metabolomic, and lipidomic profile associated with tumorigenesis in a murine model of ulcerative colitis, highlighting the risks of specific intestinal dysbiosis in genetically predisposed subjects.

**What you need to know:** *Background and context:* Colitis-associated colorectal cancer arises from complex host-environment interactions, including gut microbiome influences, driving chronic inflammation, with the intestinal lumen environment remaining a largely unexplored potential risk factor in cancer development.

*New findings:* Winnie mice in specific pathogen-free conditions developed severe colitis, and a novel juvenile colon dysplasia and cancer, with gut microbiome changes driving colitis-associated cancer initiation and progression.

*Limitations:* We identified a pro-inflammatory microbial/metabolic signature promoting colitis-to-CAC transition in Winnie mice, with FMT confirming microbiota-driven tumor susceptibility. However, further research is needed to pinpoint the key bacteria-metabolite-lipid combination driving CAC.

*Clinical research relevance:* This newly characterized microbiota-metabolome-based model of CAC, challenges the dogma of cancer as a non-transmittable disease, providing a foundation for developing microbiota-based strategies for CAC prevention and treatment.

*Basic research relevance:* Unlike genetic or chemically induced models, the Winnie mouse model uniquely serves as a dual model for spontaneous colitis and juvenile CAC, offering a fast, 100% penetrant phenotype that enhances reliability, accelerates research, and provides valuable insights into IBD and CAC.

## INTRODUCTION

The pathogenesis of colitis-associated cancer (CAC) is a multifaceted process driven by chronic inflammation, genetic and epigenetic changes, dysregulated cell proliferation, oxidative stress, and alterations in the gut microbiome. A significant risk factor for CAC is ulcerative colitis (UC), characterized by persistent inflammation of the colon and rectum, and whose prevalence has been rising, particularly in Western and newly industrialized countries^1^. UC patients with longstanding and relapsing inflammation face an approximately 2–3-fold increased risk of developing colorectal cancer (CRC)^2^. Despite advancements in surveillance and therapy, CAC remains a high-risk condition with increased mortality, often requiring colectomy as a last therapeutic option^3^.

Although the interface between chronic inflammation and carcinogenesis is an area of active investigation, the precise mechanism(s) by which some, but not all, patients with UC progress to CAC remain unknown. This complexity arises from the challenge of understanding how multiple contributing factors interact, as they are often studied independently rather than in a comprehensive and integrative manner. Emerging evidence highlights a critical role for the gut microbiome and its metabolic byproducts in modulating the development of CAC^4,5^, suggesting that microbial interactions serve as essential factors among the multiple potential drivers of CAC. The gut microbiome, a complex and dynamic ecosystem, influences intestinal homeostasis through microbial-derived metabolites that impact inflammation, immune modulation, and tumorigenesis^6^. Dysbiosis, an imbalance in microbial composition, has been implicated in promoting a pro- carcinogenic environment through mechanisms such as the production of genotoxic compounds, disruption of epithelial integrity, and induction of chronic inflammation^7–9^. Several different experimental conditions demonstrate a decrease in the development of colonic tumorigenesis under germ-free (GF) conditions, regardless of the trigger’s nature, whether genetic or chemically induced^10,11^. Thus, strategies, including fecal microbiota transplantation (FMT), probiotics, prebiotics, synbiotics, postbiotics, and dietary modifications, have been explored to enhance the effectiveness of immunotherapy by modulating the gut microbiome^12^. Furthermore, the gut microbiome has been identified as both a biomarker and a modulator of anti-tumor immunotherapy outcomes^13,14^. Indeed, combining the modulation of gut microbiome with chemotherapy has been shown to enhance treatment efficacy in CRC, suggesting that a synergistic approach to therapy may be more beneficial for these patients^15^.

Interestingly, a recent finding suggests that alterations in both the gut microbiome and its metabolic products can influence the inflammatory environment, potentially contributing to the development of CAC^16^. Additionally, metabolic pathway alterations may play a dual role in the pathogenesis of CAC. On one hand, amino acid metabolism and bile acid biosynthesis have been implicated in driving the inflammation-to-carcinoma progression characteristic of CAC^17^. On the other hand, short-chain fatty acids (SCFAs), including acetate, propionate, and butyrate, have been shown to have an inverse correlation with inflammation, suggesting a potential protective effect against both chronic inflammation and colorectal cancer (CRC) development. Several studies also demonstrate that lipids play a key role in CAC by influencing inflammation, immune responses, and tumor progression. Pro- inflammatory lipid mediators regulate inflammatory pathways associated with cancer development^19^ and are present in colon tumors, indicating their influence on the tumor microenvironment^20^. Additionally, dysregulated lipid metabolism promotes CAC progression by disrupting antitumor immune responses^21^. Overall, these findings suggest that CAC develops through a multifaceted and intricate process, also involving the gut microbiome and its related metabolome/lipidome.

To gain deeper insights into the histopathologic and morphologic changes in CAC triggered by various factors and explore potential treatments, different murine models have been developed, including chemically induced, spontaneous, genetically modified, and adoptive transfer models^5^. While these models have provided key insights into UC pathogenesis, replicating the full complexity of the human disease remains a significant challenge. Notably, the Winnie mouse strain has emerged as a promising model of UC-like colitis, closely mirroring the disease’s clinical and histopathological characteristics. Its ability to develop spontaneous colitis due to a point mutation in the *Muc2* gene, leading to epithelial barrier dysfunction, highlights the critical role of barrier damage in triggering inflammation, making it a valuable tool for studying UC progression and potential therapy^22,23^. Unlike chemically or genetically induced colitis models, Winnie mice naturally exhibit complex interactions among immune, epithelial, and neuronal cell populations and colonic microbiota, closely mirroring the complexity of human UC. Additionally, their spontaneous and fully penetrant phenotype, with disease manifesting as early as four weeks of age, provides a significant advantage over IL-10 KO and MDR1α KO mice, where colitis results from genetic deletion and is not consistently present in all individuals^5^. Notably, before this study, despite exhibiting chronic and progressive inflammation in the large intestine, Winnie mice had never been reported to develop spontaneous CAC unless crossed with *APC^Min/+^* mice.

With the intent to gain a comprehensive perspective on the impact of the intestinal microbiota and associated metabolome/lipidome in colitis-prone Winnie mice, we rederived Winnie mice, originally in conventional (CONV) housing, into a specific-pathogen-free (SPF) facility (SPF Winnie)^5,23^. Differently from what we previously published on CONV Winnie mice^24^, we observed a more severe colitis in SPF Winnie mice, even without crossing with *APC^Min/+^* mice as we did in the prior study, and surprisingly, an early onset of colonic tumorigenesis (at four weeks of age) with the extent and progression of disease increasing over time (up to 20 weeks of age). Using shotgun metagenomics and untargeted metabolomic and lipidomic approaches, we characterized the -omics profiles of the two colonies (*i.e*., SPF Winnie and parental tumor-free CONV Winnie), revealing a microenvironment enriched in pro-inflammatory microbes, metabolites, and lipids that may be critical in transitioning from colitis to CAC in the SPF Winnie. Finally, to confirm the necessity of gut microbiota and related metabolome/lipidome to CAC pathogenesis in SPF Winnie, we performed fecal material transplantation (FMT) using either SPF Winnie or wild-type (WT) littermate controls (Bl/6) mice as donors and GF Winnie as recipients. Our data showed that colitis was similarly observed in GF Winnie recipients, independent of the donor feces. At the same time, only the transfer of SPF Winnie fecal material resulted in inflammation-associated colonic tumors, underscoring the importance, and perhaps the existence, of a specific CAC-predisposing gut microbiome *milieu* in SPF Winnie mice. Therefore, in the present study, we report a previously undescribed model of CAC dependent on a non-tumor-associated genetic mutation, combined with a unique intestinal microbiome and metabolome. We provide further evidence that the composition and function of the gut microbiome play a non-redundant role in promoting colonic tumorigenesis through fecal microbiota transplantation (FMT) into germ-free (GF) recipients with a genetic factor, such as *Muc2*, which is involved in regulating gut homeostasis and colonic inflammation.

## MATERIALS AND METHODS

### Animal studies

Our investigations were performed under the ARRIVE guidelines and the relevant animal protocols, which were approved by both the Institutional Animal Care Committee of the National Institute of Gastroenterology “S. de Bellis” in Italy (Organism engaged for compliance of Animal Wellbeing, OPBA; DGSA Prot. 768/2015-PR 27/07/2015), and The Institutional Animal Care and Use Committee (IACUC) at Case Western Reserve University (protocol 2014-0158) in the USA. All the animal experiments performed in Italy were carried out according to the national guidelines of Italian Directive n. 26/2014 and approved by the Italian Animal Ethics Committee of the Ministry of Health-General Directorate of Animal Health and Veterinary Drugs. All animals were maintained in a controlled environment (20-22°C, 12h light and 12h dark cycles, and 45-55% relative humidity), housed in standard cages with corn bedding, and fed with a standard diet (Prolab® IsoPro® RMH 3000, LabDiet). Mice were sex matched for all the experiments and the experiments were performed on mice aged 4, 8, 12, 16 and 20 weeks (each timing is specified throughout the text). All mice were on C57BL/6J background (https://www.jax.org/strain/000664<u>#</u>) and all the experiments were performed on sibling littermates.

Mouse endoscopies were conducted under isoflurane anesthesia according to the IACUC protocol using a flexible uteroscope (Olympus) and following our laboratory standard procedure for imaging and scoring^25^.

### Histology analysis

Mice were euthanized according to the ARC protocols and their colons were explanted, fixed in 10% formalin for 24h, and washed with and stored in 70% EtOH before paraffin embedding, sectioning and staining for Hematoxylin & Eosin (H&E) to be scored for inflammation by a trained pathologist. Colon length and weight were measured as indicators of colonic inflammation at sacrifice. The colon/body weight index was calculated as a percentage, representing the ratio of colon weight to body weight for each mouse.

### 16S rRNA sequencing

180 mg of stool sampled from CONV and SPF 20-week-old Winnie mice were processed for DNA extraction using QiaAMP Powerfecal Pro DNA kit (Qiagen) following the manufacturer’s protocol. For tissue samples, total DNA was extracted using the Qiagen Blood and Tissue kit and processed like stool DNA. Sequencing was carried out on Illumina MiSeq platform by the CWRU Genomics core facility, with paired end method; the V3-V4 region was amplified for analysis of diversity inside the Bacteria Domain as previously done^7^.

### Deep shotgun metagenomics and functional analysis

To better characterize our mice’s microbiota, we performed a deep shotgun and functional analysis from the same samples used for the 16S rRNA analysis. CosmosID (Germantown, MD, US) carried out the analysis and bioinformatics.

DNA libraries were prepared from extracts using the Nextera XT DNA Library Preparation Kit (Illumina) and IDT Unique Dual Indexes with total DNA input of 1 ng. Libraries were then sequenced on an Illumina NovaSeq 6000 platform 2x150bp. Initial QC, adapter trimming and preprocessing of metagenomic sequencing reads are done using BBduk^26^. The quality-controlled reads are then subjected to a translated search against a comprehensive and non-redundant protein sequence database, UniRef90. The mapping of metagenomic reads to gene sequences are weighted by mapping quality, coverage and gene sequence length to estimate community wide weighted gene family abundances as described by Franzosa *et al.*^27^. Gene families are then annotated to MetaCyc^28^ reactions (Metabolic Enzymes) to reconstruct and quantify MetaCyc metabolic pathways in the community as described by Franzosa *et al.*. Furthermore, the UniRef90 gene families are also regrouped to GO terms^29^ in order to get an overview of GO functions in the community.

### Metabolomics and lipidomics sample preparation

200 mg of stool samples from the same animals used for the 16S sequencing were processed using the OMNImet®·GUT kit before the metabolite extraction. The samples were then processed and the metabolites were extracted as previously reported^30^. Following centrifugation, polar metabolites and lipids fractions were separately collected, dried by a SpeedVac (Savant, Thermo) and stored until analysis. For the assessment of repeatability and instrument stability over time, a QC strategy was applied. Metabolomics and Lipidomics were performed on two different analytical platforms^31^.

### Untargeted lipidomics and metabolomics

Lipid analysis was performed by RP-UHPLC-TIMS on a Thermo Ultimate RS 3000 coupled online to a TimsTOF Pro quadrupole Time of flight (Q-TOF) (Bruker Daltonics, Bremen, Germany). Lipidomics MS data alignment, filtering and annotation was performed with MetaboScape 2021 (Bruker). Untargeted metabolomics was carried out by HILIC-HRMS on a Thermo Ultimate RS 3000 coupled to a Q-Exactive quadrupole-Orbitrap (Thermo Scientific, Bremen, Germany). Metabolomics MS data alignment, filtering and annotation was performed with MS-DIAL v4.80. Detailed conditions for LC and MS parameters together with all pre-processing steps are reported elsewhere^32,33^. Univariate and multivariate statistics was performed with MetaboAnalyst 6.0 (https://www.metaboanalyst.ca). Data preprocessing consisted of the following steps: all metabolites or lipids missing in more than 50% in QCs and 75% in real samples were excluded, subsequently data were normalized, log transformed and auto scaled. For missing values and zeros, one-fifth of the minimum value for the target molecule in the dataset was used for replacement. A preliminary investigation was carried out using Principal Component Analysis (PCA) on pre-processed data. Significant (p-value < 0.05) metabolites and lipids and their HMDB codes were employed to build a pathway enrichment analysis by Enrichment function on MetaboAnalyst.

### Integration of -omics

We used a supervised N-integration approach to jointly analyze multi-omics datasets, including 16S rRNA microbial profiles, lipidomics, metabolomics, and functional pathways derived from metagenomic data, in the context of CONV and SPF Winnie mice. Principal coordinates were first extracted from each omics layer, followed by integrated partial least squares discriminant analysis (PLS-DA) using the mixOmics R package. This method constructs latent components by maximizing the covariation between datasets while optimizing class discrimination and integration. The model employs M-fold or leave-one-out cross-validation across a user-defined parameter grid to determine the optimal sparsity level. The resulting parameters facilitate the simultaneous selection of key discriminative features across all datasets.

### Fecal microbiota transplantation

A total of 400 mg of stool from 5 male and 5 female of 20-week-old SPF Bl/6 and Winnie mice were dissolved in 10 mL of sterile PBS to a final concentration of 40 mg/mL. 8-week-old germ-free Bl/6 and Winnie mice were gavaged with 200 µL of this stool suspension, twice a week for two weeks. Mice were monitored each day and weighed every week until the sacrifice 4 weeks after the first gavage. Mice were euthanized and their colon was explanted, measured and weighted. The colon was cut longitudinally, and tumor areas were measured before fixing the colon in formalin.

### Statistical analysis

Data analysis for dot plots and histograms was performed on GraphPad 10 software using parametric and non-parametric tests, one-way ANOVA followed by Tukey’s multiple comparison test or Kruskal-Wallis test followed by Dunn’s test as appropriate for group comparisons (see individual figure legends).

## Acknowledgments

This work was supported by National Institutes of Health Grants DK042191, DK055812, and DK091222, which were awarded to Fabio Cominelli. The authors also acknowledge the Mouse Models Core and the Histology/Imaging Core of the Cleveland Digestive Disease Research Core Center (DK097948). They thank Ashtyn Balasko for managing the mouse colonies. Finally, the authors thank the Case Western Reserve University Genomics Core Facilities and Cytometry Core Facilities for their assistance with 16s rRNA sequencing and flow cytometry, respectively.

## Author Contributions

GV and SD shared the first authorship and contributed to the acquisition, analysis, and interpretation of data, drafting of the manuscript, and statistical analysis. MC and FC are senior co-authors and contributed to the study concept and design, analysis and interpretation of data, drafting and critical revision of the manuscript for important intellectual content, attainment of funding, and study supervision. GV, SD, and BC performed the experiments. EMS, DL, LZ, FDC, FM, and VC performed the computational analysis of the data. TTP and PC contributed to the drafting and critical revision of the manuscript for important intellectual content. All authors contributed to refining the study protocol and approved the final supervision.

## RESULTS

### SPF Winnie mice show a pro-tumorigenic phenotype as compared to CONV Winnie mice

Winnie mice from a CONV facility in Italy were transferred to a SPF facility in the U.S. Heterozygote Winnie mice (*Muc2*^+/-^) were used for breeding to obtain Winnie mice (*Muc2*^-/-^) and their littermate wild-type control (*Muc2*^+/+^, Bl/6 mice) from the same breeding (Figure 1A). Weight comparisons between the original CONV and the new SPF colonies from weaning (4 weeks) to sacrifice (20 weeks) revealed significant differences (Figure 1B). While SPF Winnie mice were initially heavier at 4 weeks, their growth stalled around 8 weeks, whereas CONV Winnie mice continued gaining weight (Figure 1B). By 12, 16, and 20 weeks, CONV Winnie mice had significantly higher body weight, likely due to a worsening disease phenotype in SPF Winnie mice (Figure 1B, C). Despite no differences in colon length at sacrifice (data not shown), SPF Winnie mice had a higher colon weight-to-body weight (BW) ratio (Figure 1D), indicating colon-specific pathology. In line with that, a time course analysis of H&E-stained colon showed increased histology score over time in SPF vs. CONV Winnie mice (Figure 1E), which presented only mild inflammation even at later time points (Figure 1E). In line with the inflammatory score, tumor area borders progressively expanded over time, even at early time points (4 weeks) (Figure 1F). Unlike the previously characterized Winnie-*APC^Min/+^* model^24^, tumors in SPF Winnie mice were specifically located in the proximal and medial colon as indicated by the light box and stereomicroscopic images (Figure 2A, *left* and *right panels*, respectively). Conversely, SPF Bl/6 littermates obtained from the same SPF-Winnie breeders do not show tumors or signs of inflammation. Data collected over time demonstrate that GF Winnie and their SPF Bl/6 counterparts do not develop tumors and inflammatory phenotypes. Similarly to the anatomical characteristics of human CAC, endoscopic observations confirmed the presence of neoformations protruding into the lumen of SPF Winnie mice colon (Figure 2B, *upper row*, white-dotted area) located close to the splenic flexure, a mark of the transition from the distal to the medial part of the colon^25^. Additionally, the mucosa in tumoral areas was thicker in SPF Winnie mice, obscuring blood vessels visible in their littermate controls as well as in both GF Bl/6 and Winnie mice (Figure 2B, *lower row*). Histological analysis of 20-week-old SPF Winnie mice revealed dysplastic crypts and severe inflammation (Figure 1G, *middle image plus inset*), while CONV Winnie mice only showed diffuse inflammation (Figure 1G, *upper image*). Additional inflammatory features in SPF Winnie mice included crypt elongation, immune infiltration, and crypt abscesses; these features are absent in SPF Bl/6 and GF Winnie mice (Figure S1, *upper panel*). PAS-Alcian blue staining confirmed a general reduction of mucus-secreting cells and the loss of acid mucins (Alcian blue staining) in SPF Winnie mice compared to SPF Bl/6 and GF Winnie mice (Figure S1E). Moreover, aberrant crypts exhibited a unique PAS staining pattern, characterized by a reduced number of mucus-secreting cells along the epithelium and a concentrated magenta staining in the crypt centers, indicating a shift toward neutral mucin production during tumor progression (Figure S1, *lower panel*). Our results emphasize the role of microbiota in driving CAC in SPF Winnie mice, as reflected by increased inflammation, greater tumor burden, and distinct mucosal changes, highlighting the influence of environmental factors on disease progression.

**Figure 1:**
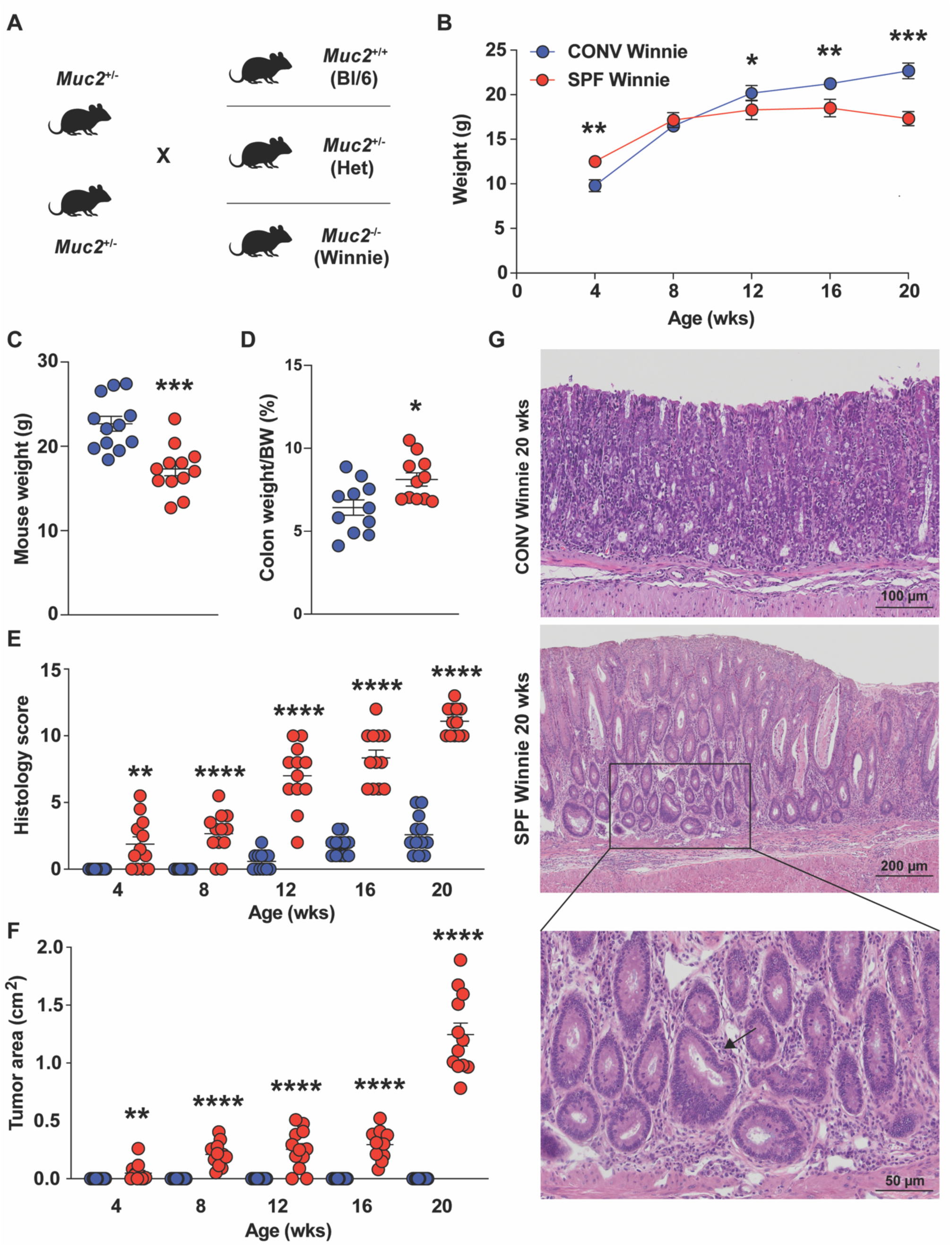
Increased disease severity and emergence of colonic tumors in SPF- vs. CONV- raised Winnie mice. **(A)** Breeding strategy used to generate Winnie mice and their littermate controls (Bl/6) in specific pathogen-free (SPF) and conventional (CONV) facilities. (**B**) Weight curve of SPF vs. CONV Winnie mice measured every 4 weeks until sacrifice at 20 weeks. (**C**-**G**) Necroscopic and histological evaluations of colonic inflammation and tumorigenesis comparing SPF vs. CONV Winnie. (**C**) Weight, and (**D**) colon weight/body weight (BW) ratio percentage of experimental mice measured at the time of sacrifice. Progression of (**E**) disease severity over time as evaluated by a GI pathologist blinded to H&E-stained slides of colon samples, using an established scoring system. (**F**) Tumor area over time was calculated by Image J using a necropsy image of the colon taken at the time of sacrifice. (**G**) Representative H&E images of CONV (*upper panel*) and SPF (*lower panel*) Winnie mice at time of sacrifice (10X magnification); inset highlights dysplastic crypt (arrow), 20X magnification. Data presented in the dot plots are expressed as mean ± SEM, with *P*-values calculated by unpaired *t*-test or Mann-Whitney; *n*=12 for each experimental group; \**P*<0.05; \*\**P*<0.005; \*\*\**P*<0005; \*\*\*\**P*<0.0001.

**Figure 2:**
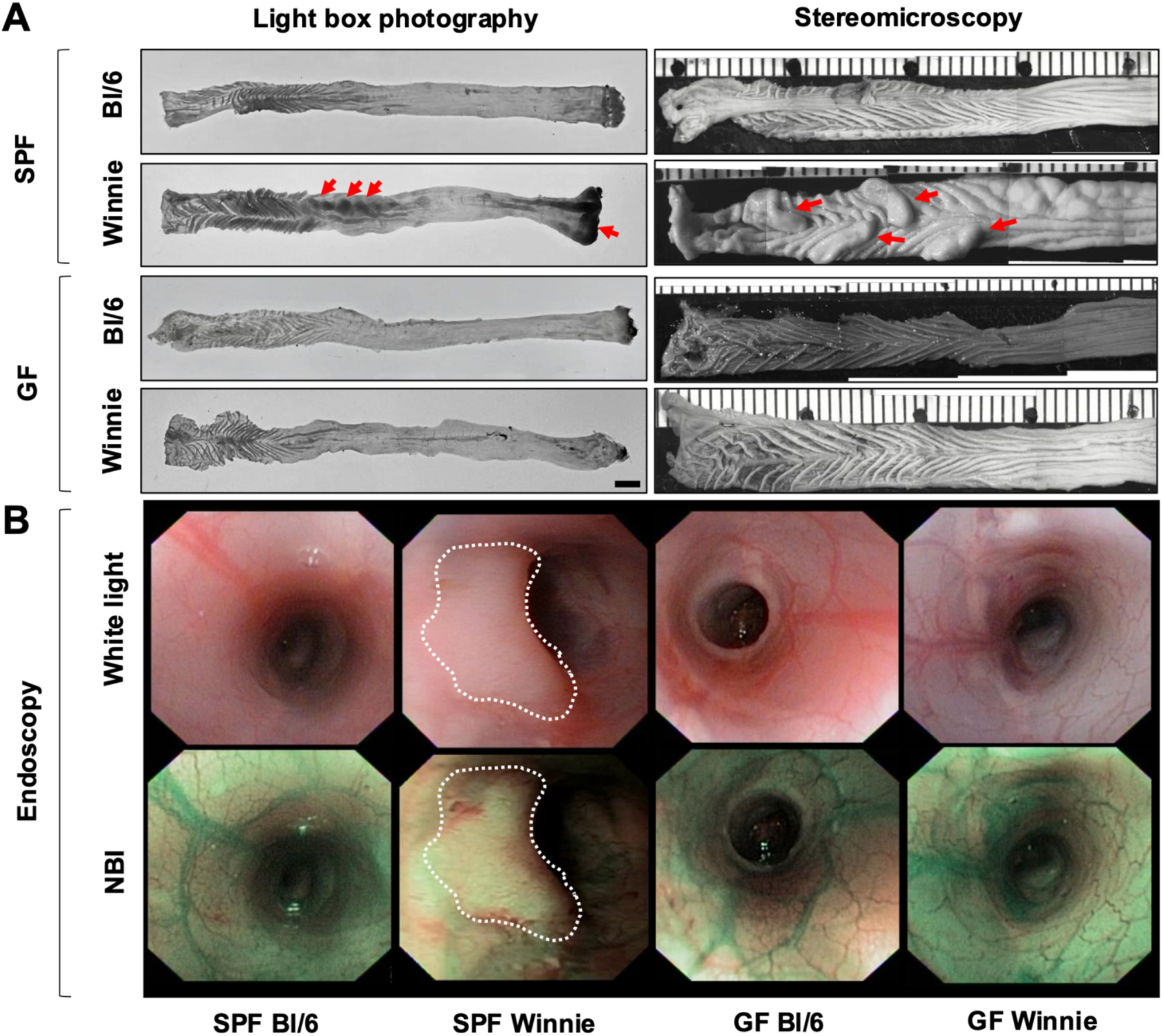
Development of colon tumors in SPF Winnie mice is dependent on the presence of gut microbiota. **(A)** Representative light box (*left panels*) and stereomicroscopic (*right panels*) images of fresh and fixed, respectively, 20-week-old SPF and germ-free (GF) wild-type (Bl/6) and Winnie colons. (**B**) Representative endoscopic images of 20-week-old SPF (*left panels*) and GF (*right panels*) Bl/6 and Winnie mice using white light (*upper panels*) and narrow band imaging (NBI) (*lower panels*). The white-dotted outline highlights the representative tumor area.

### Specific microbial populations shape the fecal and tumoral phenotype of SPF Winnie mice

Next, we performed 16S rRNA analysis to compare stool microbiota from 20-week-old SPF and CONV Winnie mice. We observed a distinct separation between SPF and CONV Winnie mice in terms of alpha diversity, as shown by observed OTU and Shannon index (Figure 3A and B, respectively), as well as beta diversity (Figure 3C). Composition of the most abundant (>1%) microbial genera from SPF and CONV Winnie mice is indicated by the pie charts shown in Figure 3D. Specifically, we found a significant upregulation of *Prevotellaceae_UCG-NK3B31*, *Faecalibaculum*, *Dubosiella*, *Lachnospiraceae_UCG-006,* and *Anaerosporobacter* genera in SPF vs. CONV Winnie, while an opposite regulation was detected for the genus *Prevotellaceae_UCG-001* (Figure 3E). We also compared tumor-associated microbiota to non-tumor mucosa without observing differences in beta diversity (Figure 3F). Looking at the most abundant microbial genera composition in this comparison (Figure 3G), four genera were found to be differentially represented in SPF Winnie. Specifically, *Gastranaerophilales*, *Rhodospirillales,* and *Lachnospiraceae_AC2044* genera were less abundant, while *Alloprevotella* was more abundant in tumors compared to non-tumor mucosa of SPF Winnie mice (Figure 3H). Overall, these results reveal specific bacterial genera, differently enriched in SPF Winnie mice, suggesting a potential link between microbiota alterations and tumor progression in SPF Winnie mice.

**Figure 3:**
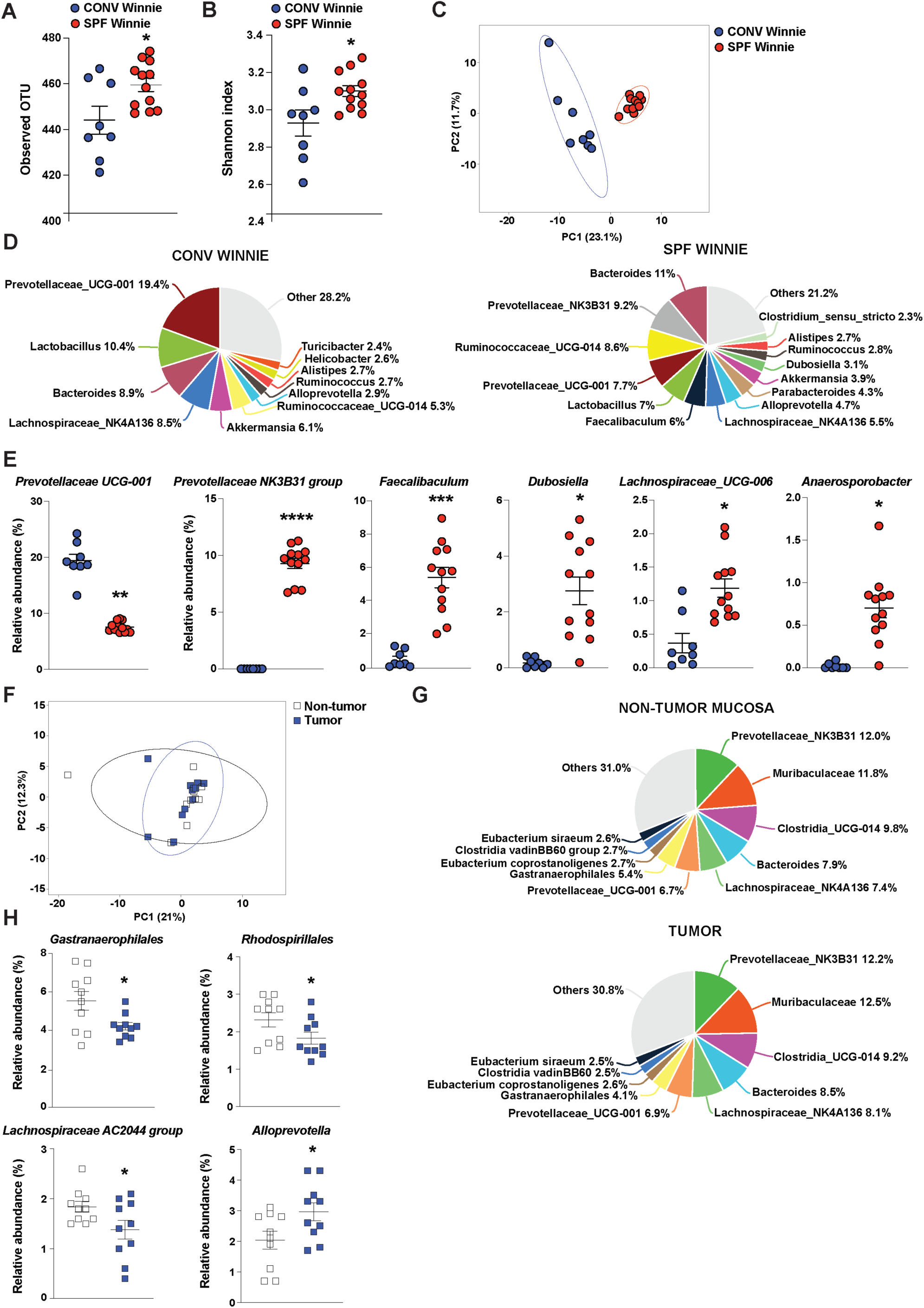
Microbiome composition differs in tumor-bearing SPF vs. CONV Winnie mice, with differential regulation of *Gastranaerophilales, Rhodospirillales, Lachnospiraceae AC2044 group*, and *Alloprevotella*. (**A**-**E**) 16S rRNA sequencing on fecal total microbiota from 20-week-old SPF (*n*=12) and CONV (*n*=8) Winnie mice. (**A**) Observed OTU (operational taxonomic unit), (**B**) Shannon diversity index, and (**C**) PCA (principal component analysis) were assessed in the two experimental groups. (**D**) Pie charts for SPF (*right*) and CONV (*left*) Winnie stool, indicating percentages of the most abundant genera (>1%). (**E**) Relative abundance (expressed as percentage), of significantly represented genera in the stool of 20-week-old SPF (*n*=12) and CONV (*n*=8) Winnie mice. (**F**-**H**) 16S rRNA sequencing on non-tumor and tumor-associated microbiota of 20-week-old SPF Winnie mice. (**F**) PCA, with (**G**) pie charts depicting non-tumor mucosa (*upper*) and tumor-associated mucosa (*lower*), showing percentages of most abundant genera (>1%), and by (**H**) relative abundance (expressed as percentage) of significantly represented genera in SPF Winnie. Data presented in the dot plots are expressed as mean ± SEM, with P-values calculated by unpaired t-test or Mann-Whitney; n=12 for each experimental group; \**P*<0.05; \*\**P*<0.005; \*\*\**P*<0.0005; \*\*\*\**P*<0.0001.

### Stool metabolomics and lipidomics highlight altered sphingolipid and nucleotide metabolism in SPF Winnie mice

We conducted metabolomic and lipidomic analyses on stool samples from 20-week-old SPF and CONV Winnie mice. Principal Component Analysis (PCA) revealed two distinct clusters corresponding to the experimental groups (Figure 4A). The volcano plot identified 300 metabolites and lipids significantly enriched (Figure 4B, *blue*) and 76 decreased (Figure 4B, *grey*) in SPF compared to CONV Winnie mice. Enrichment analysis of the top 25 metabolites linked them to inflammatory bowel disease (IBD), inflammation, cancer, and autoimmune disorders (Figure 4C), with further associations to nucleotide metabolism and bioenergetic pathways (Figure 4D). The heatmap confirmed the distinct separation of the top 50 significant metabolites and lipids in fecal samples from the two groups (Figure 4E). Supplementary Figure S2 presents dot plots of some of the most significant metabolites and lipids. The dot plots in Fig. 4C-D report the overview of the enriched metabolite sets, where the size of the dots per metabolite set represents the Enrichment Ratio and the color intensity represents the p-value. Among the major lipid classes differentially abundant between SPF and CONV Winnie mice, sphingolipids -particularly ceramides (CerS), hexosylceramides (HexCerS), and sphingomyelins (SMs)-were the most modulated subclasses. SPF mice exhibited higher fecal levels of SMs, HexCerS, and CerS. Additionally, lysophosphatidylcholines (LPCs) and lysophosphatidylethanolamines (LPEs) were significantly more abundant in SPF feces. Regarding the polar metabolome, we observed a pronounced alteration in purine and pyrimidine nucleotide metabolism. The levels of AMP, CMP, GMP, and UMP were all elevated in SPF mice, suggesting an upregulation of nucleotide metabolism, which is often associated with tumor growth. Consistently, NAD+ levels were also increased in SPF mice, reflecting higher metabolic demands, as cancer cells require significantly more NAD+ than normal cells to sustain their elevated energy consumption.

**Figure 4:**
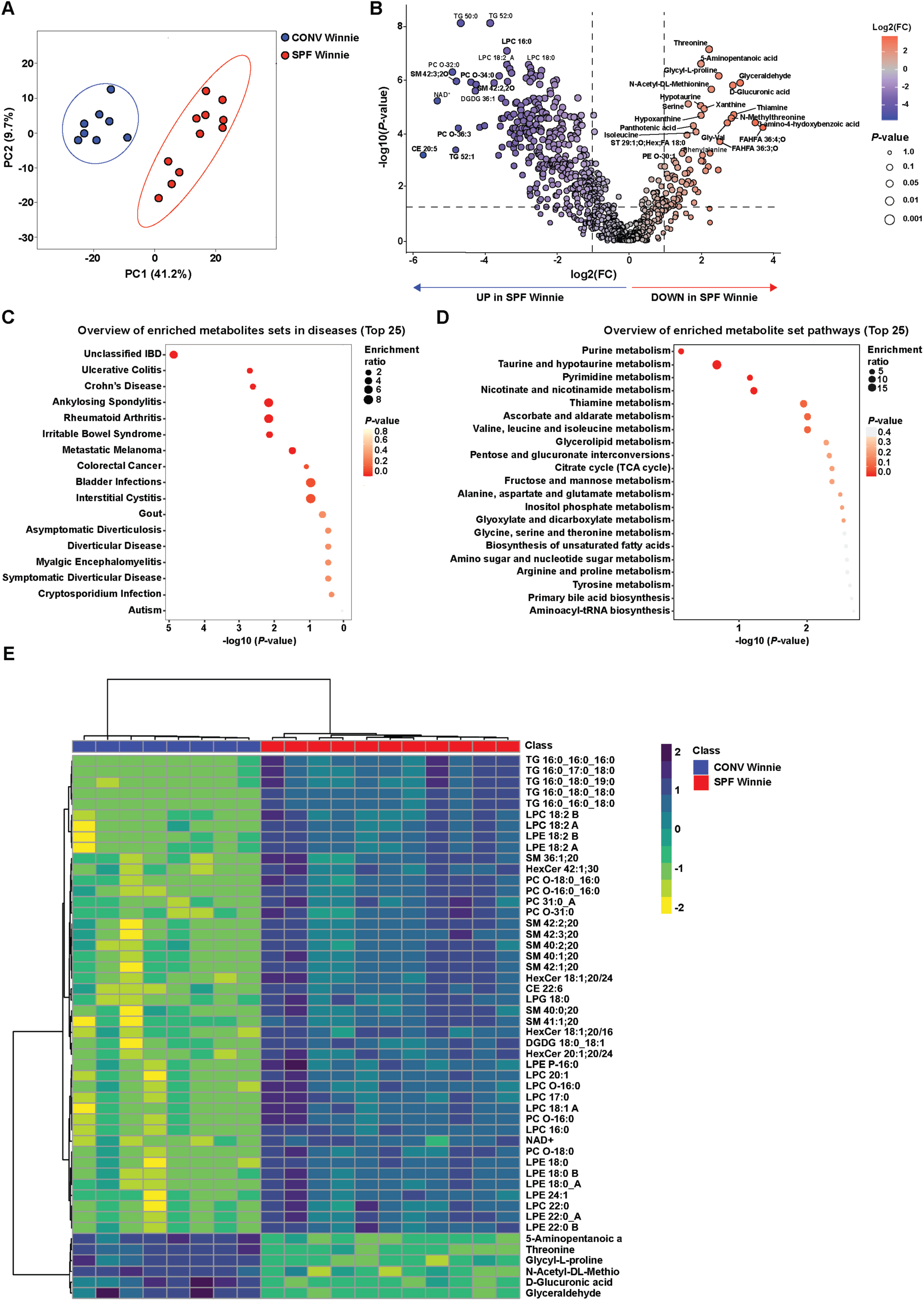
Metabolipidomic profiling of stool samples from 20-week-old SPF and CONV Winnie mice reveals distinct metabolomic and lipidomic alterations. Metabolipidomics of stool from 20-wk old CONV vs. SPF Winnie mice shown by (**A**) Score plot of PCA analysis relative to metabo-lipidomics, (**B**) volcano plot indicating significantly increased (*blue*) and decreased (*red*) metabolites and lipids in SPF Winnie mice. (**C**) Quantitative enrichment analysis overview showing top 25 related diseases, (**D**) metabolic pathways ranked according to *P*-value (0.0 equals <0.05) and fold enrichment, and (**E**) heatmap for the top 50 significant metabolites and lipids of SPF relative to CONV Winnie mice.

### Shotgun metagenomic analysis reveals distinct microbial profiles and metabolic pathways in SPF and CONV Winnie mice

To build on the insights gained from 16S sequencing, we performed shotgun sequencing on 20-week-old SPF and CONV Winnie mice. This analysis also revealed a clear distinction between the two experimental groups, as demonstrated by the score plot of PCA (Figure 5A) and microbial species clustering in the heatmap (Figure 5B). Despite these differences, a close phylogenetic relationship was observed among the most abundant and differentially represented species in SPF and CONV Winnie mice (Figure 5C). Examining the significantly regulated microbial species, we found that *Duncaniella muris*, *Muribaculum intestinale*, *Parabacteroides distasonis*, *Phocaeicola sartorii*, *Lactobacillus taiwanensis,* and *Faecalibaculum rodentium* were more abundant in CONV Winnie stool samples (Figure 5D). Interestingly, tumor-associated species such as *Alistipes finegoldii*, *Phocaeicola vulgatus,* and *Bacteroides fragilis* were enriched in SPF Winnie mice. Additionally, in SPF Winnie mice, *Akkermansia muciniphila* and *Ligilactobacillus murinus* were significantly more abundant, even though these two species are frequently indicated as protective for colitis and cancer. Next, we performed a functional analysis of these bacterial species, revealing an enrichment of proinflammatory and iron-related pathways in the SPF group (Figure 5E). These pathways correlated with the bacterial species that were more abundant in SPF Winnie mice. On the contrary, SCFA metabolism was more strongly associated with bacteria characteristic of CONV Winnie mice (Figure 5F). Shotgun analysis further indicated that SPF-associated bacteria exhibited heightened iron-capturing and metabolic activity, with increased representation of iron ion binding, heme binding, and iron homeostasis pathways. Additionally, several of the most prominent pathways aligned with the elevated metabolites detected in our Winnie cohorts, including those involved in thiamine, pantoate, nucleotide, and fatty acid metabolism.

**Figure 5:**
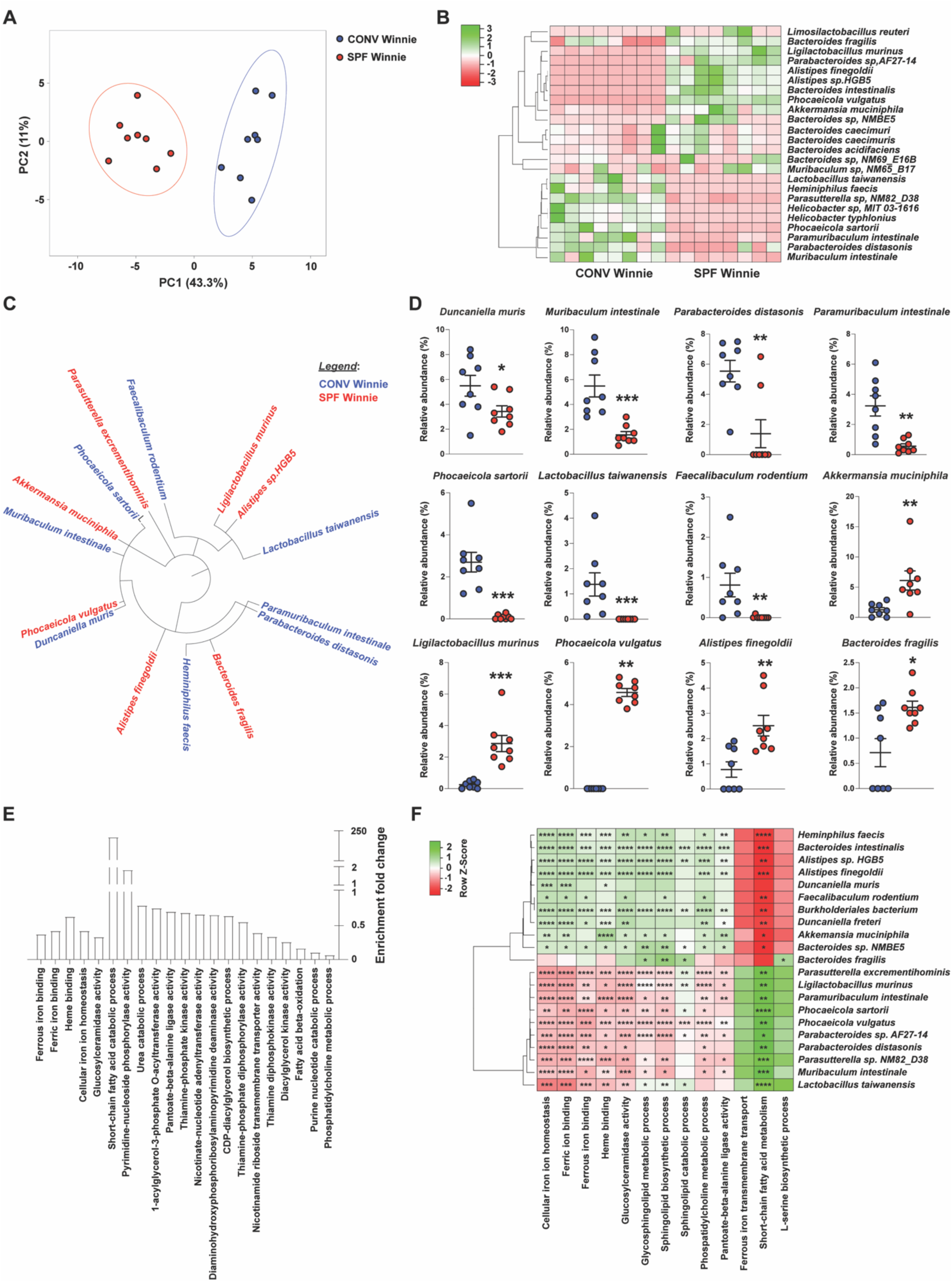
Shotgun analysis identifies bacterial strains and related pathways driving a pro- tumorigenic phenotype in SPF vs. CONV Winnie. Fecal samples from 20-week-old SPF vs. CONV Winnie mice were analyzed and presented by (**A**) PCA, (**B**) heatmap of the most abundant bacterial species (>1%), (**C**) circular dendrogram highlighting significantly different bacterial species, and (**D**) dot plots showing relative abundance (expressed as percentage) of the most abundant species. (**E**) Enrichment fold-change of specific pathways relevant in colitis and colitis-associated cancer, comparing fecal material from SPF and CONV Winnie mice. (**F**) Spearman’s correlation between bacterial species and relevant pathways enriched in SPF vs. CONV Winnie mice. Data presented in the dot plots are expressed as mean ± SEM, with *P*-values calculated by unpaired *t*-test or Mann-Whitney; *n*=8 for each experimental group; \**P*<0.05; \*\**P*<0.005; \*\*\**P*<0005; \*\*\*\**P*<0.0001.

### Multi-omic correlation analysis reveals distinct microbial and metabolic profiles driving CAC in SPF Winnie Mice

The use of the same fecal sample for 16S rRNA sequencing, metabolomics/lipidomics, and shotgun sequencing analysis enabled a direct comparison across datasets, allowing us to uncover meaningful relationships between microbial composition, metabolic profiles, and lipidomic changes. Specifically, we performed a multi-omic correlation analysis that confirmed a distinct segregation between SPF and CONV Winnie mice, consistent with the patterns observed in individual analyses, when all datasets were combined (Figure 6A). A two-dimensional projection further validated this separation, reflecting the clustering trends identified in the independent analyses (Figure 6B). Focusing on dimension 1, the average distances between the principal coordinates are noticeably larger in the SPF group compared to the CONV group. Moreover, we observed strong positive correlations between differentially abundant bacterial genera, metabolites, and lipids across the groups (Figure 6C). The differential contributions of specific genera, lipids, and metabolites to the SPF and CONV Winnie phenotypes are shown in Figure 6D, suggesting distinct microbial and metabolic signatures associated with each condition. This indicates a potential functional relationship between microbial composition and metabolic alterations in SPF Winnie mice. These findings offer a comprehensive, systems-level perspective on the microbial and metabolic shifts in SPF Winnie mice, underscoring the intricate interplay between gut microbiota and host metabolism in CAC development in SPF Winnie mice.

**Figure 6:**
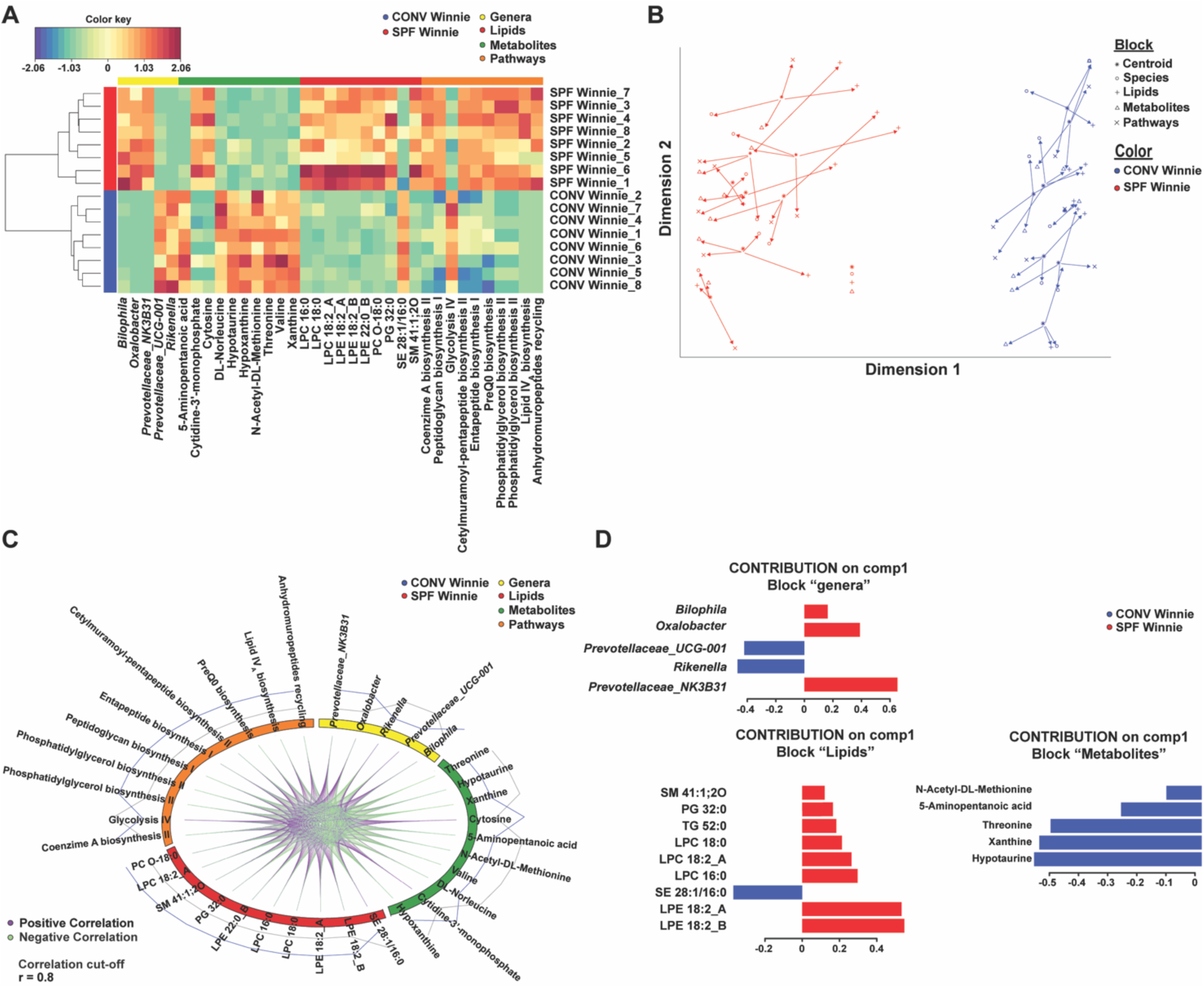
Integration of multi-omic analyses shows strong correlation between metabolomics, lipidomics, and bacterial genera. The heatmap represents the correlation structure extracted from the four omics datasets (**A**). The correlation of each original feature pair is determined by each of their correlations with the components from the integrative method. The arrow plots depict the similarities and discrepancies between a given sample across the four datasets, which can be seen (refer to the legend) (**B**). Short arrows indicate strong agreement between datasets, while long arrows highlight significant disagreement. Each sample is represented by a centroid and associated with four data types. The circos plot depicts the correlations between each feature of each dataset. The top selected features of each dataset are shown. Lines are only drawn for correlations above 0.8 (cutoff = 0.8) to reduce visual clutter (**C**). The Loading generated from the PLS-DA applied to the multiomic data highlights the variables that discriminate between treatment conditions. Blue indicates the SPF group, while red represents the CONV group (**D**). *n*=8 for each experimental group.

### FMT from SPF Winnie but not SPF Bl/6 mice, specifically increases dysplasia index in GF Winnie mice

To confirm the role of microbiota in CAC onset in SPF Winnie mice, we performed FMT by transferring stool from 20-week-old SPF Winnie or its age-matched parental control (SPF Bl/6) into GF Winnie mice. GF Bl/6 recipient mice served as negative controls (Figure 7A). Specifically, FMT was administered by gavage (40 mg/mL) four times over two weeks, and mice were sacrificed 4 weeks after the first gavage (Figure 7B). Weight monitoring revealed a trend of weight reduction in recipients of SPF Winnie stool, regardless of genotype (GF Winnie or GF Bl/6) (Figure 7C). At sacrifice, GF Winnie mice that received SPF Winnie stool exhibited a higher colon weight/BW ratio compared to those transplanted with SPF Bl/6 stool (Figure 7D). Endoscopic analysis showed an increased score in GF Winnie mice transplanted with both SPF Winnie and SPF Bl/6 stools, though no significant difference was observed between these groups (Figure 7E). Furthermore, histological analysis identified dysplastic lesions in GF Winnie mice transplanted with both SPF Bl/6 and SPF Winnie stools, though the lesions were significantly larger in those receiving SPF Winnie stools (Figure 7F). Of note, while colonic inflammatory score did not differ between GF Winnie transplanted with SPF Winnie and SPF Bl/6 stools (Figure 7G), the dysplasia index was significantly higher in GF Winnie mice transplanted with SPF Winnie stool as compared to GF Winnie transplanted with stools from SPF Bl/6 (Figure 7H). In line with these results, GF Bl/6 mice receiving stool from either SPF Winnie or SPF Bl/6 showed no weight reduction (Figure 7C), no changes in the colon weight/BW ratio (Figure 7D), and a null score for endoscopy, inflammation, and dysplasia index (Figures 7E, G, H). These findings suggest that the microbiota from SPF Winnie mice plays a crucial role in promoting CAC development in a genetically susceptible host.

**Figure 7:**
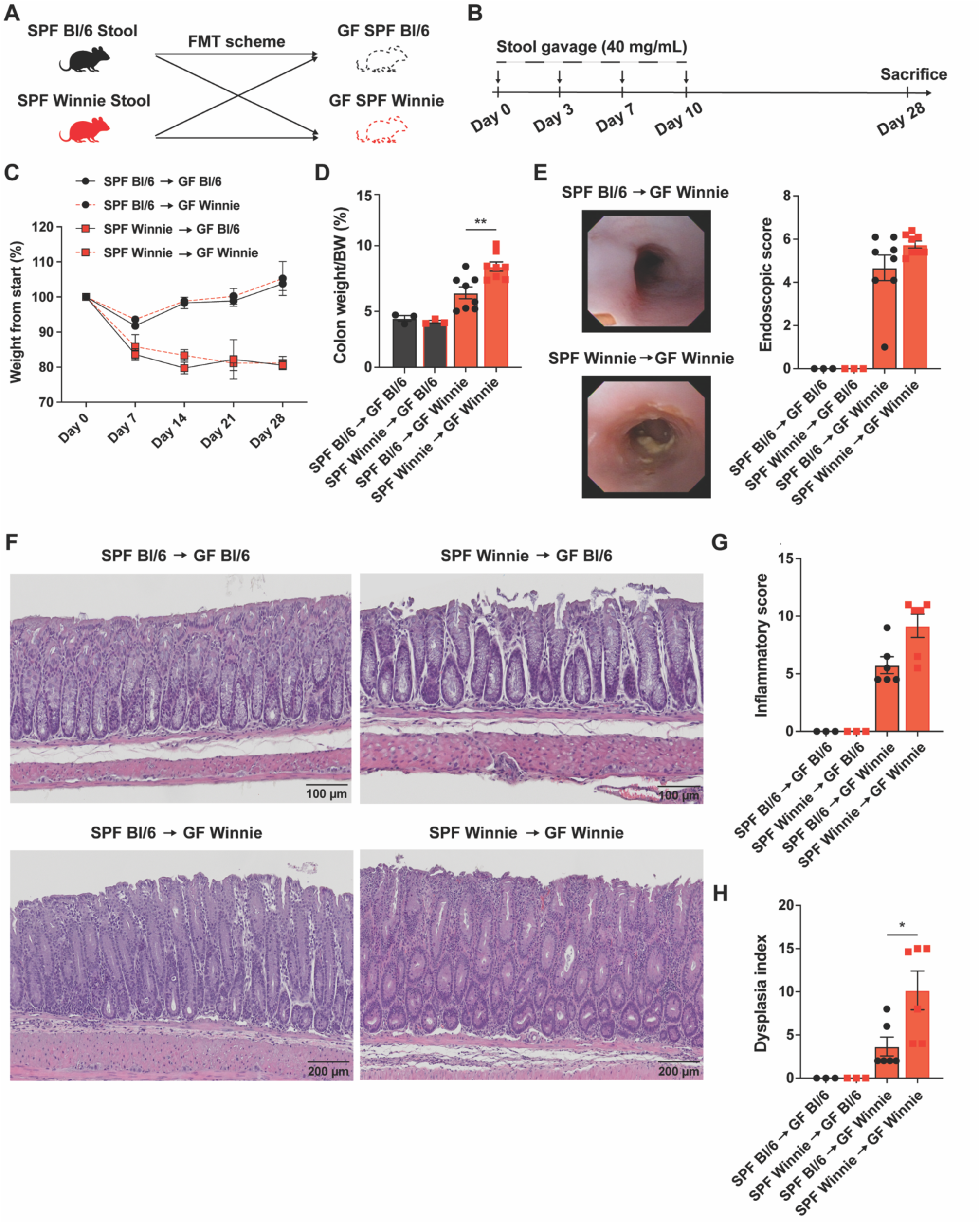
Fecal microbiota transfer (FMT) using SPF Winnie and Bl/6 donors indicates that tumor development depends on Winnie, not Bl/6 the host recipients. Schematic of fecal microbiota transplantation (FMT) experimental design (**A**) and timeline (**B**) using SPF Winnie and Bl/6 mice donors into GF Winnie and Bl/6 recipient mice. (**C**) Changes in weight of experimental mice over time, and (**D**) colon weight/BW ratio (expressed as percentage) measured at sacrifice. (**E**) Representative white light endoscopic images (*left panels*) of GF Winnie recipient mice receiving donor SPF Winnie (*lower panel*) or SPF Bl/6 (*upper panel*), with associated endoscopic scores (*right panel*). (**F**) Representative H&E images of experimental groups (10X magnification), with associated histological assessment of (**G**) inflammation, and (**H**) dysplasia, as evaluated by a GI pathologist blinded to H&E-stained slides of colon samples, using an established scoring system. Data presented in the histograms are expressed as mean ± SEM, with *P*-values calculated by ordinary one-way ANOVA followed by Tukey’s multiple comparison or Kruskal-Wallis test followed by Dunn’s test. Stool homogenates are pooled from *n*=4 donor mice, with *n*=3-8 recipient mice for each experimental group; \*\**P*<0.005; \*\*\**P*<0005.

## DISCUSSION

Here, we described a novel model of colitis-associated cancer (CAC) with early onset that is dependent on the gut microbiome and its metabolites. Following rederivation into an SPF facility, Winnie mice exhibited a more severe colitis phenotype and, notably, spontaneous CAC as early as four weeks of age. In contrast, CONV Winnie mice developed only mild colitis as previously reported in other facilities^5^, with no overt signs of tumorigenesis. We demonstrated the essential role of the gut microbiome by observing that GF Winnie mice were protected from colitis and colon tumor development. Using shotgun metagenomics, metabolomics, and lipidomics, we characterized a distinct pro-inflammatory microbial and metabolic signature that potentially drives the transition from colitis to CAC. FMT using either SPF Winnie or WT (Bl/6) donors into GF Winnie recipients demonstrated that while colitis developed regardless of donor, only FMT from SPF Winnie donors resulted in CAC, revealing a critical microbiota-driven, host-specific susceptibility to tumorigenesis in this model.

Compared to CONV Winnie mice, SPF Winnie mice showed a decreased body weight but increased colon weight. This was mainly due to increased inflammation and a larger tumor area. Indeed, SPF Winnie colon presented tumors in the proximal and medial parts, a feature resembling what is often observed in the ascending and transverse portions of CAC patients^34,35^. Furthermore, the macroscopic features of these tumors are comparable to those of flat and serrated adenomas typical of CAC^36^. The dysplastic lesions had an early onset, consistent with the incidence of CAC in IBD patients^37^, unlike sporadic CRC or previously established models of CAC, like AOM/DSS or Winnie/*APC^min/+^*, where tumors appear as polyps and have a late onset in patients^38^.

Microscopically, SPF Winnie mice show increasing signs of inflammation in their colon, with dysplastic lesions appearing in 4-week-old mice. Lesions that grew slowly and steadily with the age of the mice, similar to those in CAC patients^39^, showed a steep increase at 20 weeks of age, concurrent with a higher dysplasia grade. On the other hand, CONV Winnie did not develop any tumors but only mild inflammation. Thus, the different microenvironment and the differences in the housing facility led to distinct phenotypes; the increased inflammation and the formation of dysplastic lesions might be caused by the SPF facility being cleaner than the CONV one, hence creating perturbations in the intestinal microbiota.

One of the most striking aspects of our model is the unique microbiome composition we identified through 16S rRNA sequencing. Despite alpha diversity being almost the same, beta diversity changed dramatically, as evidenced by the clear separation of the two Winnie populations on the PCA plot; moreover, several bacterial genera were differently represented in the stool of these mice. Within the *Bacteroidetes* phylum, opposite abundance was noticed for *Prevotellaceae-UCG001* and *Prevotellaceae NK3N31*. The first was significantly less represented in the SPF Winnie microbiota, while the second was absent in CONV mice and more than 9% in the SPF Winnie. Both groups are associated with fiber consumption and SCFA production; however, *Prevotellaceae-UCG001* has recently been linked to the inhibition of colorectal cancer progression in FMT experiments. *Faecalibaculum*, significantly more represented in the SPF-Winnie fecal material, is a lactic acid producer with a disputed role in colon cancer. It is possible that the abundance of lactic acid in the lumen affects luminal pH, alters gut metabolism, and promotes the tumor microenvironment^40^. *Faecalibaculum* has been suggested as a marker of colorectal cancer (CRC). Nonetheless, its role in SCFA production may support the idea of a pro-homeostatic switch to pathogenic under particular conditions^41^. Similarly, other SCFA-producing groups, known for their anti-inflammatory and tolerogenic potential^42–44^, including *Dubosiella*, *Lachnospiraceae UCG-006,* and *Anaerosporobacter*, are increased in the SPF-Winnie fecal material. The hampered immune response may indeed accelerate tumor growth.

When we looked at the tumor-associated microbiota, we noticed that *Alloprevotella* was significantly higher in CAC tissue than the paired normal tissue of SPF Winnie, similar to what was previously reported in humans^45^. *Gastranaerophilales*, *Lachnospiraceae AC2044,* and *Rhodospirillales* were reduced considerably in the healthy tissue of SPF Winnie mice, but their functional role is not fully elucidated. These bacteria may be associated with dysplastic mucosa or part of the mucosal biofilm that promotes tumor growth.

We performed stool metabolomics and lipidomics on the same samples to gain a deeper insight into microbial activity. Similarly to what was observed for the 16S sequencing, samples from the SPF and CONV colonies separated well on the PCA plot. We obtained several more abundant metabolites in SPF Winnie mice, related to the pathways of energy production and amino acid recycling. Among the metabolites related to nucleotide metabolism, adenosine 3’-monophosphate and adenosine 3′,5′-cyclic monophosphate were the most detected in SPF Winnie, confirming what was previously reported in a fecal samples metabolomics analysis from CRC patients^47^. There is evidence that most of the metabolites we detected are produced in areas of ongoing inflammation, as immune cells utilize them to sustain their activity. Next, with stool lipidomics, we observed an interesting perturbation in the sphingomyelin/ceramide pathway. Sphingomyelin is a membrane phospholipid that can be hydrolyzed by sphingomyelinases (SMases) to generate ceramide^48^. SPF Winnie mice had increased sphingomyelins in contrast to CONV mice, which had more ceramides in their stool^49^. *Bacteroidetes* rise non-significantly in the SPF Winnie and are the only sphingolipid producers in the gut microbiome^50^. Using Sptlc2 KO mice, Li et al. demonstrated that the block of the ceramide *de novo* synthesis pathway dramatically affected the mucus layer and E-Cadherin expression in the gut, suggesting a direct link between ceramide and gut barrier function^51^. LPC, which we found increased in SPF Winnie’s stool, is often linked to increased inflammation and colorectal cancer. Bacterial sphingolipids are indeed associated with increased inflammation and cellular division, which is connected to the onset of cancer; conversely, ceramides have been shown to exhibit anti-inflammatory potential. Moreover, other metabolites we found increased in CONV Winnie stool, such as the sterols ST 29:1 and ST 28:1 (stigmasterol and its derivatives), are linked to decreased inflammation and protection from DSS colitis^52^.

To confirm that stool bacteria produce these metabolites, we performed a functional analysis on the data obtained from the shotgun metagenomics. What we found was indeed a more active sphingomyelin biosynthetic pathway in SPF Winnie stool compared to CONV mice. Furthermore, we observed a significant increase in the SCFA metabolism pathway in CONV Winnie stool, indicating that some bacteria in their microbiota are actively producing SCFAs with anti-inflammatory effects. Moreover, SPF bacteria exhibited increased iron metabolism, a sign often correlated with more pathogenic species, which can also increase the risk of cancer formation in IBD conditions. The presence of tumor-related and tumor-promoting bacteria was also observed with the shotgun analysis, where species like *Alistipes finegoldii*^53^ and *Bacteroides fragilis*^54–56^ were more abundant in SPF Winnie mice; the same can be said with the *Phocaeicola*^57^ species and *Ligilactobacillus murinus*^58,59^, while CONV mice had increased abundance of *Duncaniella muris* and *Lactobacillus taiwanensis* that are supposed to be protective against sustained inflammation^60,61^. Recent evidence suggests that *the presence of Akkermansia muciniphila* in the gut microbiome may be a signature of CAC susceptibility. Indeed, Winnie mice exhibit increased *Akkermansia* abundance compared to Bl/6 littermates, which may initiate tumorigenesis in the presence of other specific bacterial species. The abundance of *Akkermansia muciniphila* might also indicate a more labile mucus layer that favors the epithelial exposure to luminal antigens and inflammatory mediators.

With a comprehensive bioinformatic approach, we combined 16S, shotgun metagenomics, metabolomics, and lipidomics data, and this cross-analysis confirmed all the single inputs we had. Moreover, we employed an integrative bioinformatic approach, which has been recently used to characterize patients with early-onset CRC^64^. We obtained a correlation signature of bacteria, metabolic pathways, and metabolites that is representative of the CONV and SPF Winnie phenotypes. Surprisingly, *Akkermansia’s* central role in cancer overlaps with our correlation. To further confirm the microbial signature of the SPF Winnie phenotype, we performed FMT experiments using GF Winnie mice. We transplanted the whole SPF Winnie stool into GF Winnie and Bl/6 recipients. GF Winnie recipients exhibited colonic inflammation and dysplasia, whereas SPF Bl/6 microbiota significantly mitigated this outcome. Our model stands out as a suitable recipient of FMT, exhibiting consistent microbial engraftment and replicating the donors’ phenotype well, which is in line with previous results. The stool microbiota was responsible for the formation of dysplastic lesions, which resembled those observed macroscopically and microscopically in the stool donors. Despite these encouraging results, further studies are necessary to determine whether a specific group of bacteria induces this phenotype or if it is a result of a larger bacterial community. Previous results by our group lean towards the second hypothesis, biofilm and bacteria are indeed responsible for increased intestinal inflammation, thus the SPF tumorigenesis is likely to originate by a network of bacteria species that is absent in CONV mice and that exploits “good” bacteria metabolites and their immunosuppressive abilities to trigger intestinal inflammation and cancer onset^66,67^.

Overall, our findings reveal an intricate relationship between specific microbial communities and metabolic profiles that differ significantly from established models^68,24,9^, suggesting that this new model can provide new insights into the pathophysiology of CAC.

In conclusion, the present data demonstrate that environmental exposure may result in CAC in susceptible subjects. These results underscore the non-redundant roles of the microbiome and metabolism in the pathogenesis of CAC. The involvement of the gut microbiome in cancer has been speculated^69^. Still, to the best of our knowledge, these are the first results to demonstrate a direct connection between FMT and CAC development in a mouse model that has never been described as susceptible to colon cancer development. The unique microbiome and metabolic profiles observed in the two different facilities not only enhance our understanding of disease mechanisms but also open avenues for innovative therapeutic strategies. As we continue to unravel the complexities of the microbiome-cancer nexus, our findings underscore the importance of integrating microbiological and metabolic research into cancer studies, potentially leading to more effective prevention and treatment strategies for patients with CAC.

---

**Figure S1:**
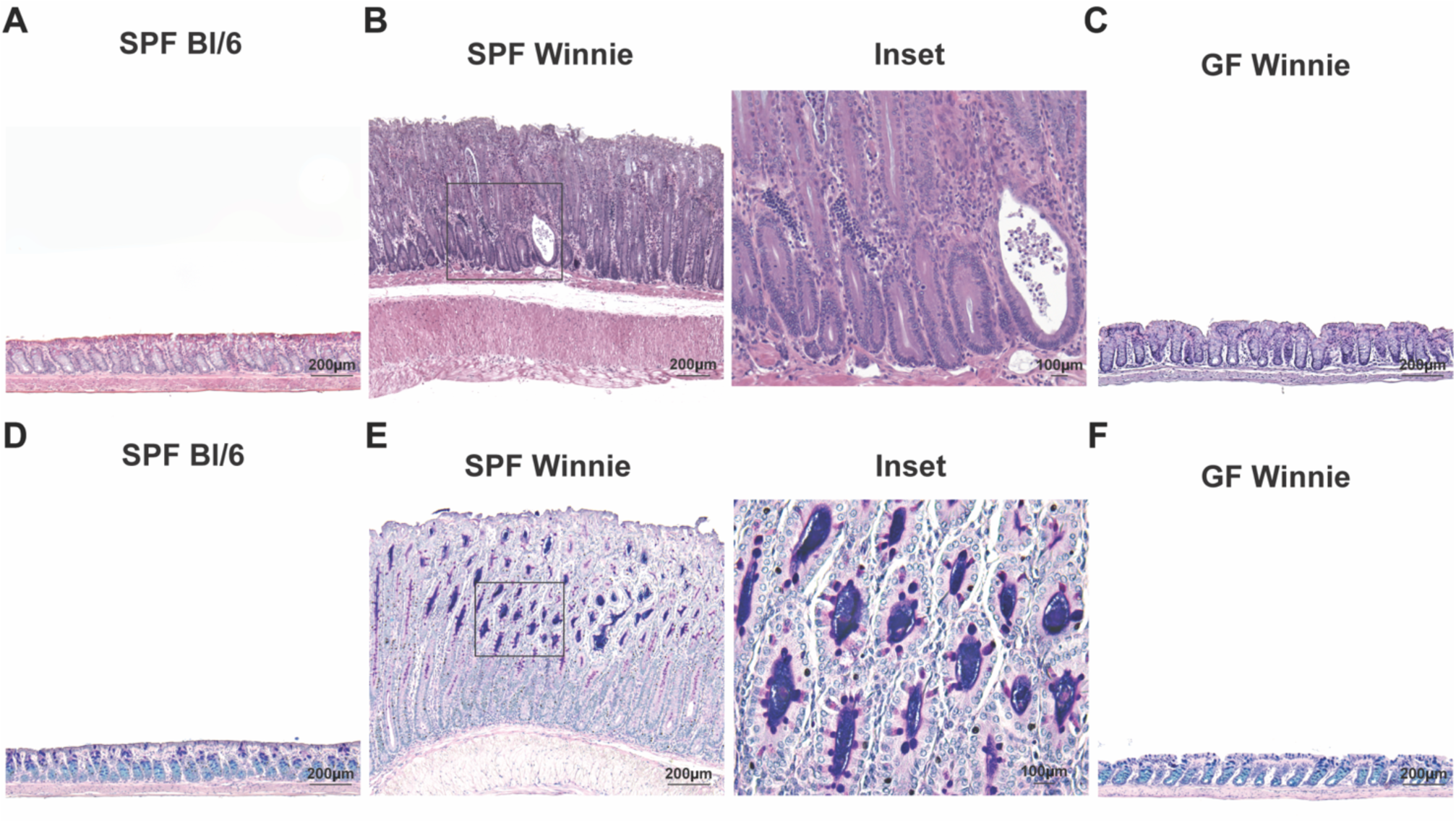
(**A**-**C**) Representative H&E images of colon tissue from 20-week-old SPF Bl/6 (**A**), SPF Winnie (**B**), and GF Winnie (**C**) mice; inset from SPF Winnie mice highlights aberrant crypts, inflammatory infiltrates, and crypt abscesses. Magnification: 5X+2, except for SPF Winnie 5X+1.65 and its inset: 20X+1.25. (**D**-**F**) Representative images of PAS/Alcian-stained colon from 20-week-old SPF Bl/6 (**D**), SPF Winnie (**E**), and GF Winnie (**F**) mice; inset highlights low staining for acid mucins (*light blue*) and an increase of neutral mucins (*magenta*) in aberrant crypts of 20-week-old SPF Winnie mice. Magnification: 5X+2, except for SPF Winnie inset: 20X+2.

**Figure S2:**
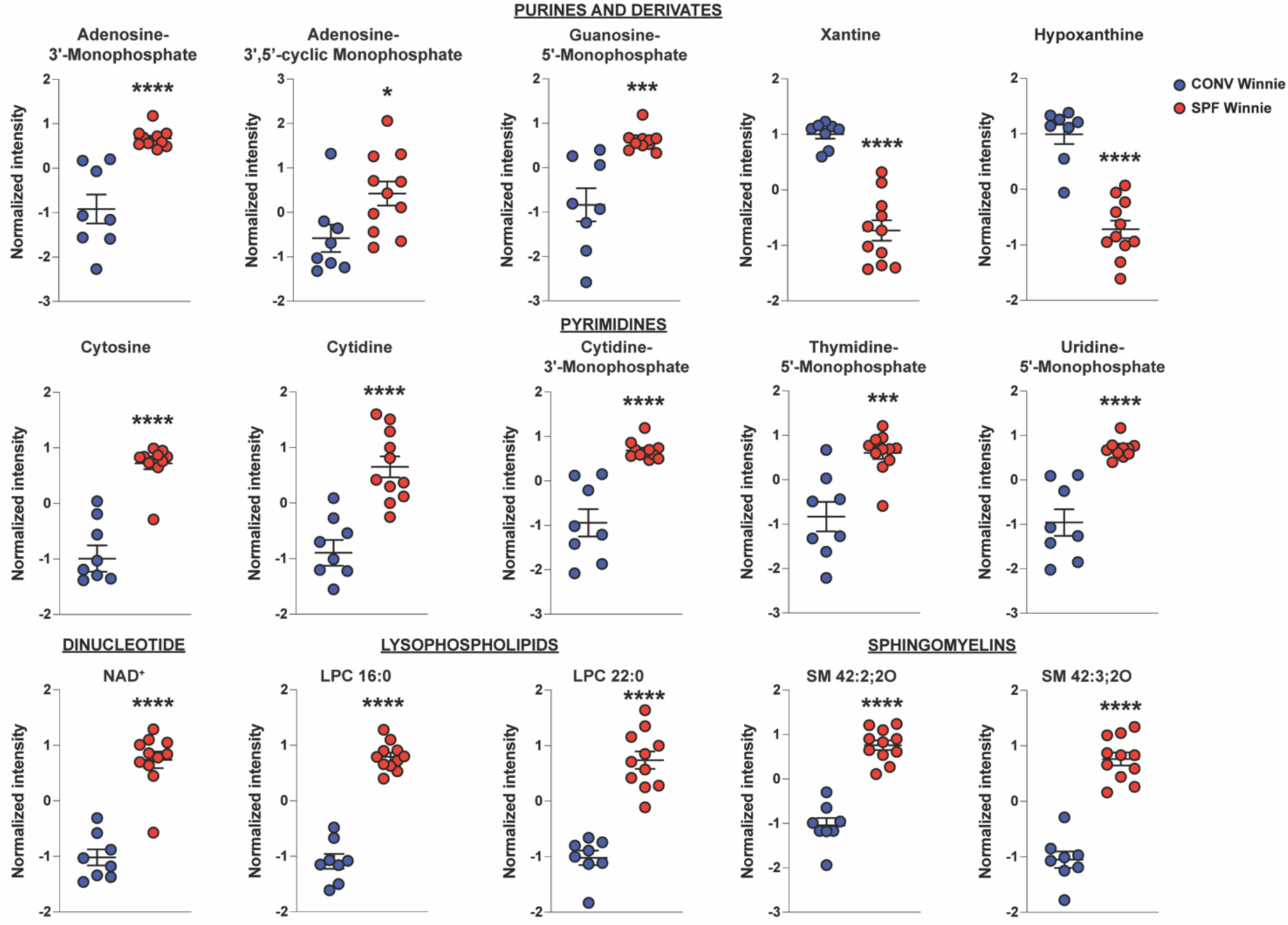
Metabolipidomic profiles differ in tumor-bearing SPF vs. CONV Winnie mice, with alterations in purine and derivatives, pyrimidines, dinucleotide, lysophospholipids, and sphingomyelins. Dot plots for normalized intensity of highly significant metabolites and lipids found in 20-week-old SPF (*red*) vs. CONV (*blue*) Winnie stool divided by compound class, *i.e*., purine and derivatives, pyrimidines, dinucleotide, lysophospholipids, sphingomyelins. Data presented in the dot plots are expressed as mean ± SEM, with *P*-values calculated by unpaired *t*-test or Mann-Whitney; *n*=12 (SPF mice), *n*=8 (CONV mice); \**P*<0.05; \*\*\**P*<0.0005; \*\*\*\**P*<0.0001.

